# Quantitative proteome dynamics across embryogenesis in a model chordate

**DOI:** 10.1101/2023.10.04.559613

**Authors:** Alexander N. Frese, Andrea Mariossi, Michael S. Levine, Martin Wühr

## Abstract

The evolution of gene expression programs underlying the development of vertebrates remains poorly characterized. Here, we present a comprehensive proteome atlas of the model chordate *Ciona*, covering eight developmental stages and ∼7,000 translated genes, accompanied by a multi-omics analysis of co-evolution with the vertebrate *Xenopus*. Quantitative proteome comparisons argue against the widely held hourglass model, based solely on transcriptomic profiles, whereby peak conservation is observed during mid-developmental stages. Our analysis reveals maximal divergence at these stages, particularly gastrulation and neurulation. Together, our work provides a valuable resource for evaluating conservation and divergence of multi-omics profiles underlying the diversification of vertebrates.

## Introduction

Urochordates are the nearest extant relatives to vertebrates and share several morphological and genomic traits (Delsuc et al. 2006). Their comparatively simple development and physiology make them attractive models to study the evolutionary origins of mechanisms in vertebrates, which have often evolved to higher complexity (Abitua et al. 2015, 2012; Berthelot et al. 2018; Christmas et al. 2023). Research on the model urochordate *Ciona* has substantially advanced the field of evolutionary and developmental biology, illuminating conserved mechanisms for body plan establishment, cellular organization, and gene regulatory networks (Abitua et al. 2015, 2012; Stolfi et al. 2010, 2015; Horie et al. 2018). These discoveries underscore the value of employing the relatively simple *Ciona* model to study processes in more complex organisms, including humans.

The assembly of the *Ciona* genome (Dehal et al. 2002) represented a significant landmark, triggering a swift surge in genome-wide analyses that included genomics, transcriptomics, epigenomics, and more recently, single-cell studies (Cao et al. 2019; Keller et al. 2016; Kubo et al. 2010; Madgwick et al. 2019; Suzuki et al. 2016; Zhang et al. 2020; Sladitschek et al. 2020). Transcriptomic studies are typically conducted to use mRNA dynamics as a surrogate for protein dynamics, the macromolecule that conveys function. Although direct measurement of protein dynamics would be more advantageous, proteome studies have been scarce, likely due to the persistent technical challenges associated with these investigations (Lopez et al. 2017; Nomura et al. 2009a; Yamada et al. 2009; Saxena et al. 2013). Technological advancements in quantitative proteomics, including multiplexing, enable sensitive and high precision measurements of protein abundances (Pappireddi et al. 2019; Johnson et al. 2021a; Thompson et al. 2003; Demichev et al. 2020; Ammar et al. 2023; McAlister et al. 2014). Proteomics studies of early development demonstrated a weak correlation between mRNA and protein levels, suggesting the importance of post-transcriptional regulatory mechanisms (Peshkin et al. 2015a; Ghaemmaghami et al. 2003; Schwanhäusser et al. 2011; Abdulghani et al. 2019; Ghazalpour et al. 2011; Vogel and Marcotte 2012; Smits et al. 2014a).

To date, proteome regulation in development remains poorly understood particularly in evolutionary contexts. Haeckel proposed that embryonic development recapitulates evolutionary trajectories (Haeckel 1866). While this theory is no longer accepted, more recent studies investigating the phylotypic period across species have generated conflicting results when based solely on transcriptome data and when limited to short phylogenetic distances within vertebrates or long distances spanning invertebrates to vertebrates (Chan et al. 2021; Uesaka et al. 2022). While some studies support the existence of a mid-embryonic period with maximal similarity in gene expression, in line with the hourglass model, others propose an inverted hourglass model, highlighting higher conservation of gene expression in the early and late developmental phases (Levin et al. 2016a; Marlétaz et al. 2018a; Hu et al. 2017a; Schrimpf et al. 2009a; Laurent et al. 2010a). To address these limitations, further investigations utilizing proteomic approaches in comparative analyses, particularly in chordates, are needed (Gil-Gálvez et al. 2022; Marlétaz et al. 2018b; Touceda-Suárez et al. 2020).

To bridge this gap, we utilized state-of-the-art proteomics to quantify the maternal proteins deposited in unfertilized eggs, as well as proteome dynamics throughout *Ciona* embryogenesis. We then integrated this data with corresponding transcriptome information and carried out an inter-species comparison, between *Ciona* and *Xenopus*, an African clawed frog, observing evolutionary conservation and divergence of protein and mRNA dynamics during development.

## Results and Discussion

### Adapting proteomics for the analysis of *Ciona* eggs and embryos

Mass spectrometry-based proteomics is a versatile tool for studying biological systems containing proteins, although new model systems often require method adaptations. Key areas needing adaptation include sample preparation and the reference proteome. Analyzing eggs and early embryos is often challenging due to the high yolk content. For instance, in *Xenopus*, yolk makes up ∼90% of egg protein content, limiting proteomics analysis depth (Goldberger 1980). Researchers usually remove yolk through centrifugation after egg or embryo lysis (Wühr et al. 2014; Gupta et al. 2018; Sonnett et al. 2018a; Baxi et al. 2018). However, when we analyzed *Ciona* egg lysates via Coomassie-stained gels, we found no exceptionally dominant protein band (Supplementary Fig. 1a), allowing us to analyze *Ciona* lysates without yolk removal by mass spectrometry. Another concern is a high-quality protein reference database. For widely used models like humans, mice, or yeast, this is derived from the genome. However, the quality of the genome for non-canonical model organisms is often poor and a better reference database can be generated based on mRNA-seq data (Wühr et al. 2014; Evans et al. 2012).

To evaluate the quality of the *Ciona* genome for proteomics analysis, we assembled a reference proteome using publicly available RNA-seq datasets (Supplementary Fig. 1b) (Wühr et al. 2014). When using this RNA-seq based reference database, it clearly outperformed the Uniprot and KY genome (Satou et al. 2019), but only increased peptide coverage by 5% compared to the recent KY21 proteome (Satou et al. 2021) (Supplementary Fig. 1c). We decided to accept the modest decrease in identified peptides for the ease of annotation offered by the genome assembly and proceeded to use the KY21 proteome as a reference for the remainder of this study. Further examination of peptides identified only by our genome-free database reveal mis-annotated gene coding sequences, mispositioned intercistronic regions, and discrepancies in selenoprotein sequences present in the KY21 proteome (Supplementary Fig. 1d,e) (Santesmasses et al. 2017; Tsagkogeorga et al. 2012a; Satou et al. 2006). We believe that our analysis will help improve the accuracy and completeness of the *Ciona* genome annotation.

Together, our analysis reveals that the sequenced *Ciona* genome, combined with the characteristics of its eggs and embryos, is highly suitable for proteomics studies, supporting *Ciona*’s potential as a valuable model system for investigating evolutionarily conserved mechanisms among chordates.

### Absolute abundance measurement in the unfertilized egg

Using the KY21 reference database, we were able to analyze the proteome of *Ciona* eggs and embryos. First, we estimated the absolute abundance of the proteins in the *Ciona* egg. Maternally deposited proteins in vertebrate eggs play a crucial role in fertilization and shape embryonic development (Wan et al. 2008; Messerschmidt et al. 2012; Bultman et al. 2006). These proteins include a range of components, including metabolic factors, transcription factors (TFs), and signaling molecules (SMs), acting as organizers and guides to ensure the proper formation of body axes and structures. We quantified 6,102 genes, after collapsing protein isoforms (Fig. 1a, Supplementary Table 1); as expected, the most abundant protein is vitellogenin (yolk protein) followed by ATP synthase subunits, actin, and a 60S ribosome subunit (Nomura et al. 2009b). Overall, our analysis spans ∼8 orders of magnitude covering 95 TFs and 46 SMs (Fig. 1b,c). The median protein concentration is 22 nM (Fig. 1c), while the median concentrations of TFs and SMs is comparatively lower, at 5.4 nM and 3.5 nM, respectively. Among the TFs identified in the egg, are known maternal factors Gata.a, Prd-B/Prdtun2, and Zinc Finger (C2H2)-33 (Imai et al. 2004a). Among the SMs are known maternal factors β-catenin, Eph.a/Eph1, Eph.b/Eph2, Raf/Raf1, and Tll/Tolloid, Notch and Numb (Imai et al. 2004b; Ahn and Kim 2012; Walton et al. 2006). Finally, we investigated whether different subunits within the same protein complex were found at expected stoichiometric ratios. To this end, we mapped the proteins identified in the egg to known stable complexes from the CORUM database (Fig. 1d) (Tsitsiridis et al. 2022). While proteomics is comparatively weak in estimating absolute concentrations, the similar value derived from subunits of the same protein complexes suggests that our study provides reliable information for absolute protein abundances in the *Ciona* egg, which can act as a valuable resource of maternally deposited proteins for *Ciona* researchers.

**Figure 1:**
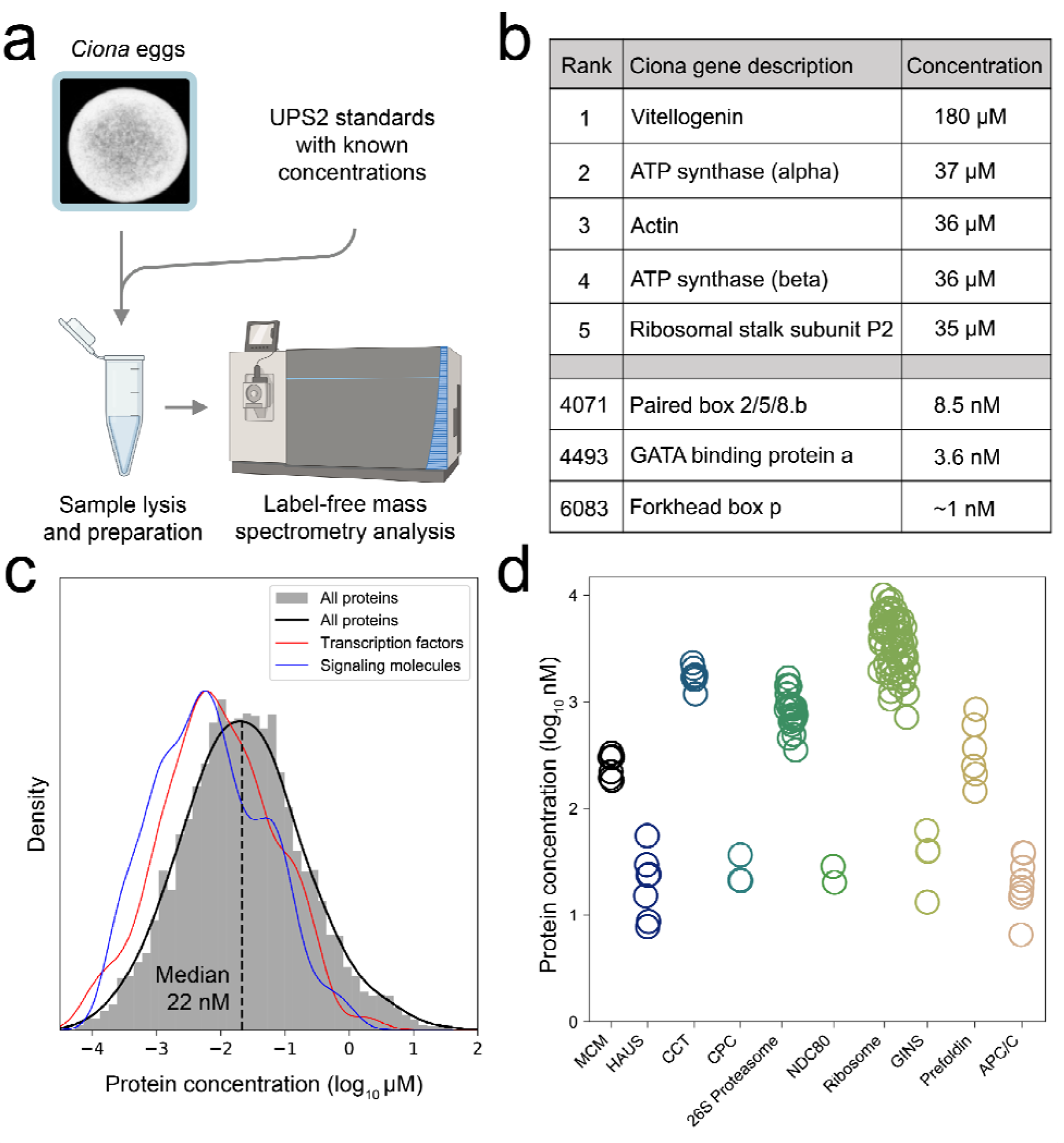
Absolute proteomics of the Ciona egg. a) Schematic of label-free proteomics utilized to determine absolute protein concentrations. Unfertilized *Ciona* eggs were lysed, and proteins of known concentrations (UPS2) were added to the lysate as a reference standard. Following normalization as outlined in the materials and methods, we were able to estimate protein concentrations for ∼6,000 proteins.b) Displayed are the five most abundant proteins, which include yolk (vitellogenin), and three known transcription factors (TFs). The complete data is provided in Supplemental Table 1.c) Histogram of all quantified proteins in the egg (gray). Kernel density estimates (KDE) of TFs (red) and signaling molecules (blue) reveal that these have a similar distribution with a lower median concentration than the global egg proteome.d) Stoichiometries of protein complexes. Estimated concentrations of subunits from a shared protein complex display similar values.

### A high-quality multi-omics atlas of *Ciona* development

Next, we measured the change of protein and mRNA abundances while the egg developed into a swimming larva. To this end, we combined accurate multiplexed proteome analysis (TMTproC) (Johnson et al. 2021b), with RNA-seq on matching samples at eight key time points spanning early embryonic development, including the maternal/zygotic transition, gastrulation, neurulation, tail elongation, and hatching of swimming larvae (Supplementary Table 2). These proteomics datasets contain 7,095 proteins (with a median of 5 peptides per protein) (Fig. 2a,b). The quantified proteins cover 38% of protein-coding genes annotated in the latest *Ciona* genome and account for approximately 50% of the expressed genes detected in RNA-seq analysis (Satou et al. 2022). This is a substantial increase compared with the only previously published proteome of *Ciona* embryogenesis which identified 695 proteins over three sampled stages using two-dimensional gel electrophoresis and MALDI-TOF/MS (Nomura et al. 2009a).

**Figure 2:**
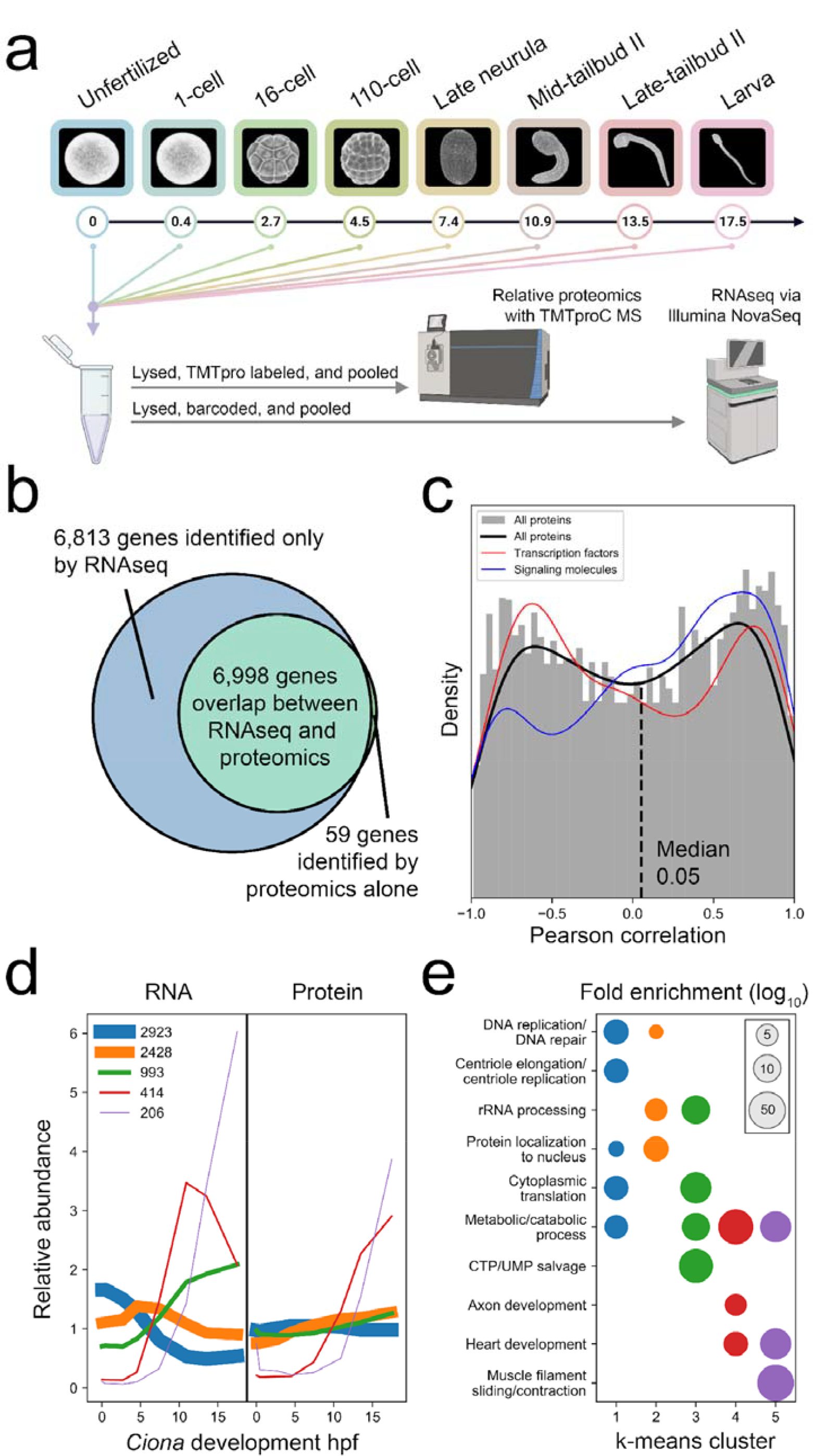
Proteome and RNA analyses during *Ciona* embryogenesis. a) Diagram of the executed proteomics and RNA-seq time-course experiments. *Ciona* embryos were harvested at eight developmental stages. Time indicates hours post-fertilization (hpf) at 18 °C. Embryos were lysed, and their mRNA and proteomes were examined through multiplexed proteomics (TMTproC) and RNA-seq.b) Venn diagram depicting genes quantified as proteins and mRNA.c) Histogram of Pearson correlations between RNA and corresponding protein dynamics throughout *Ciona* development (gray). The lines represent kernel density estimates (KDE) for all genes (black), transcription factors (red), and signaling molecules (blue). Notably, mRNA dynamics correlate poorly with protein dynamics.d) k-means clustering used to classify RNA and protein dynamics for each gene during *Ciona* development. The proteome displays greater stability than the transcriptome. The thickness of the lines scales with the number of proteins represented in each cluster, as indicated in the legend.e)GO term analysis used to discern the functional relevance of each of the clusters (indicated by matching colors) identified in d.

To assess the relationship between protein and transcript dynamics during development, we identified 6,998 corresponding genes from a combined transcript and protein quantification (Fig. 2b, Supplementary Table 3). We found that the overall Pearson correlation between mRNA and corresponding protein dynamics was very poor (median r = 0.05), consistent with previous reports (Fig. 2c) (Peshkin et al. 2015b; Smits et al. 2014b; Wegler et al. 2019; Edfors et al. 2016). This trend includes TFs with known relevance to early *Ciona* development (Supplementary Figure 2) (Imai et al. 2006).

Using k-means co-clustering of mRNA and protein dynamics, we identified 5 distinct clusters of gene dynamics (Fig. 2d). We found that the genes involved in DNA replication/repair, centriole elongation/replication, rRNA processing, and protein localization to the nucleus have maternally loaded RNA and the most static protein dynamics. Metabolic processes broadly span all of the clusters, implying that metabolic processes are not categorized by a specific dynamic pattern. Axon development, heart development, and muscle filament sliding/contraction genes are expressed at the transcript and protein level during the tailbud and larval stages of development. These data suggest that the genes in the more dynamic clusters were preferentially associated with organogenesis while the genes in the less dynamic clusters tended to drive housekeeping, or cell cycle functions (Fig. 2e).

In our study, we profiled the *Ciona* proteome and transcriptome across eight key developmental stages, spanning early embryogenesis to swimming larvae. In doing so, we generated a comprehensive proteomic atlas in *Ciona* covering 7,095 proteins with matching mRNA dynamics for nearly all proteins. This resource offers a starting point to further explore the dynamics of the protein and RNA relationship along the chordate lineage.

### Comparative analysis of proteomic dynamics in *Ciona* and *Xenopus* embryogenesis

The simple chordate *Ciona* can offer valuable insights into the distinctive developmental strategies and evolutionary novelties acquired within the vertebrate lineage. With this in mind, we compared the proteome of *Ciona* development with that of a higher vertebrate (Fig. 3a). We focused on the African clawed frog *Xenopus laevis*, which is very attractive for proteomics analysis (Sun et al. 2016; Sonnett et al. 2018a; Gupta et al. 2018; Baxi et al. 2021) resulting in probably the best characterized vertebrate proteome throughout embryogenesis. *Xenopus* and *Ciona* diverged approximately 500-600 million years ago (Hu et al. 2017b; Delsuc et al. 2018).

**Figure 3:**
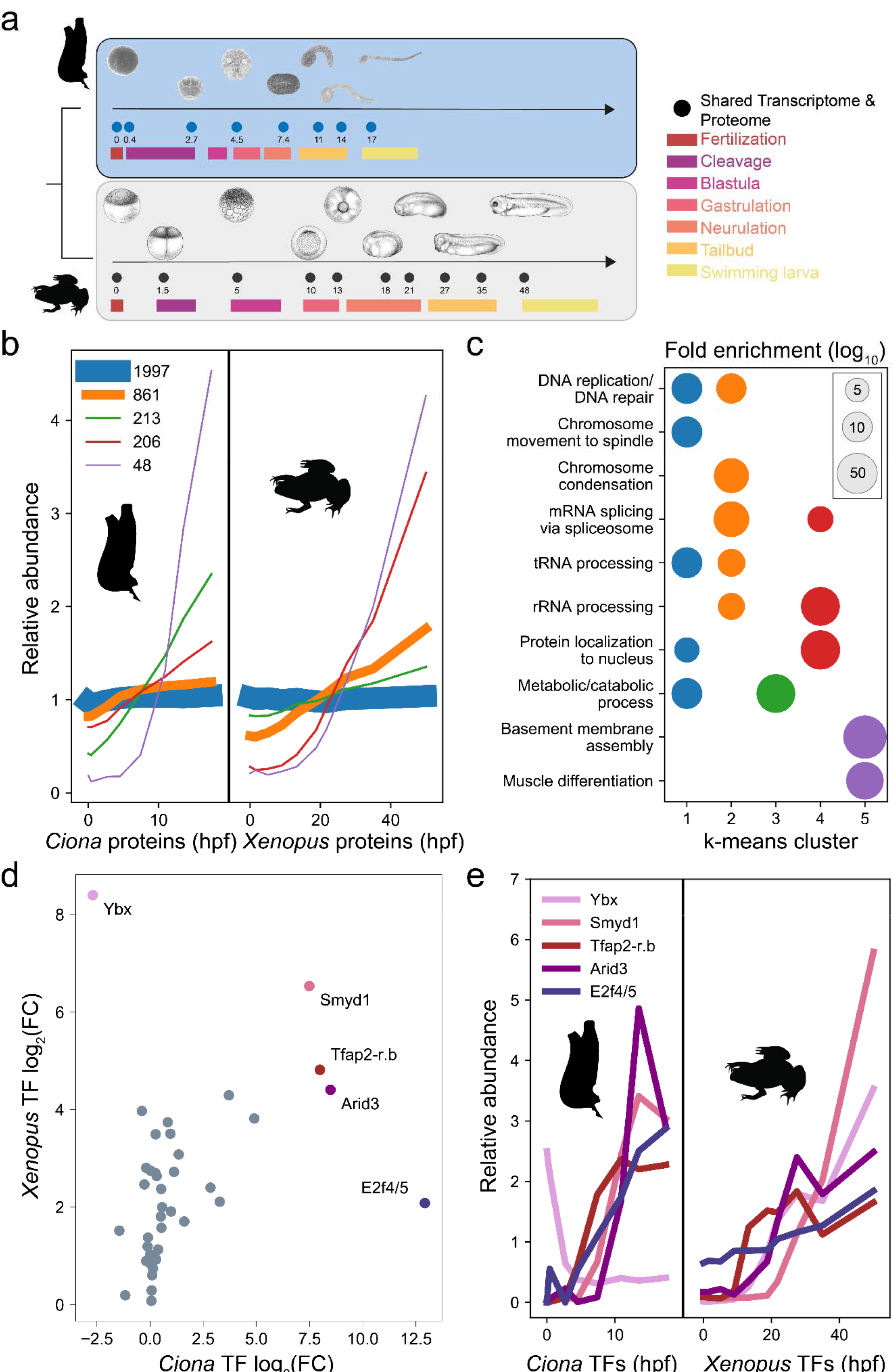
Comparison of development between chordate and vertebrate. a) Experimental design of the expanded comparative developmental transcriptome and proteome time courses. Full circles highlight stages of development sampled for RNA-seq and MS.b) k-means co-clustering of the dynamics of orthologs (3,325) between *Ciona* and *Xenopus laevis* development. The thickness of the lines scales with the number of proteins represented in each cluster. The number of proteins in each cluster are quantified in the legend.c) GO term analysis identifying the functional significance of each of the clusters from A. The color of the clusters in A is kept consistent.d) The log2 fold change (FC) protein correlation between *Ciona* and *Xenopus* TFs. Here, FC is defined as the ratio of relative protein abundance in the larva stage compared to the egg. Most transcription factors show similar behavior with the notable exception of Ybx d) The relative protein dynamics of transcription factors Ybx, Smyd1, Tfap2-r.b, Arid3, and E2f4/5. Each exhibit large fold changes in both organisms. Colors are preserved in these five proteins from the plotting in C. These TFs are canonically important for organism development by regulating transcriptional activation during the cell cycle, early muscle development, ectoderm development, gene activation through chromatin remodeling, and Nodal signaling respectively. e) Ybx exhibits signs of being maternally deposited in *Ciona*, but not in *Xenopus*, suggesting functional evolutionary divergence of this ortholog from chordate to vertebrate.

Using k-means clustering analysis, we separated the common 3,325 orthologous proteins into 5 distinct clusters, identifying significant similarities in proteome dynamics between these two species (Fig. 3b, Supplementary Table 4). More than half of the shared proteins are stably expressed in both species throughout development (Fig. 3c). This cluster is enriched for proteins involved in DNA replication, spindle formation, and chromosome movements. Clusters that capture the activity of genes involved in rRNA processing, tRNA processing, and mRNA splicing via the spliceosome show an increase in expression throughout embryogenesis in both organisms (Fig. 3b,c). Genes involved in metabolic and catabolic processes also shared an increase in expression throughout embryogenesis in both organisms, however with a more pronounced increase in *Ciona* (Fig. 3b,c). Basement membrane assembly and muscle differentiation genes have similarly high expression throughout embryogenesis in both organisms (Fig. 3b,c), including those known to have roles in late development such as Lamα5 and Smyd1 (Veeman et al. 2007; Izzi et al. 2013). These results highlight the similarities of orthologous protein dynamics during the development of these highly divergent species.

Next, our focus shifted to studying TFs during development. We looked at the relative expression of these proteins in larvae over their relative expression levels in the eggs of each organism (Fig. 3d). Overall, TFs that showed the strongest expression changes in *Ciona* tended to also increase their expression in *Xenopus*. Notably, Smyd1, Tfap2-r.b, and Arid3, which are known transcriptional regulators of muscle, ectoderm/neural crest development, and chromatin remodeling, respectively, exhibited similar patterns of expression in both species (Fig. 3e). Importantly, we observed TFs that showed different expression dynamics between the two species. The Y-box binding protein, Ybx, exhibited inverse behavior between the two organisms. In *Ciona*, Ybx appears to be maternally deposited (Fig. 3e), whereas in *Xenopus*, it is strictly expressed only after fertilization. Ybx is a highly conserved protein involved in transcriptional regulation and is a component of messenger ribonucleoprotein complexes. Understanding the underlying reasons for the differential behavior of Ybx in *Ciona* and *Xenopus* requires further investigation.

In brief, we have identified conserved and unique protein dynamics across *Ciona* and *Xenopus* through comparison of 3,325 orthologous proteins. Overall, we find strikingly high conservation of protein dynamics between the two organisms even though they are separated by ∼600 million years of evolution. This analysis therefore presents an exciting opportunity to shed light on conserved regulatory processes in chordate development, as well as the evolution of vertebrate-specific novelties.

### Proteomic comparison between *Ciona* and *Xenopus* suggests an anti-hourglass model for chordate embryogenesis with maximal diversity at mid-development

According to previous transcriptome comparative studies, gene expression is most similar during the phylotypic phase of vertebrate development, supporting an hourglass model. However, other studies support an anti-hourglass model, whereby embryos are maximally diverse at mid-developmental stages. Of course, mRNAs typically convey their functions via proteins and, to our knowledge, so far, no studies have compared protein dynamics across species and investigated the extent of diversification as a function of developmental progression. We compared mRNA and protein dynamics for each developmental stage in *Ciona* with each stage in *Xenopus* using only one-to-one orthologues.

Figure 4a shows the resulting Pearson correlation matrices for mRNA and protein dynamics. Encouragingly, the highest degree of correlation for each *Ciona* stage nearly perfectly follows the known mapping of equivalent stages for the two datasets across embryonic development. This analysis re-confirmed the presence of a bottleneck during neurulation, which represents a critical transition in vertebrate embryos (Fig 4a).

**Figure 4:**
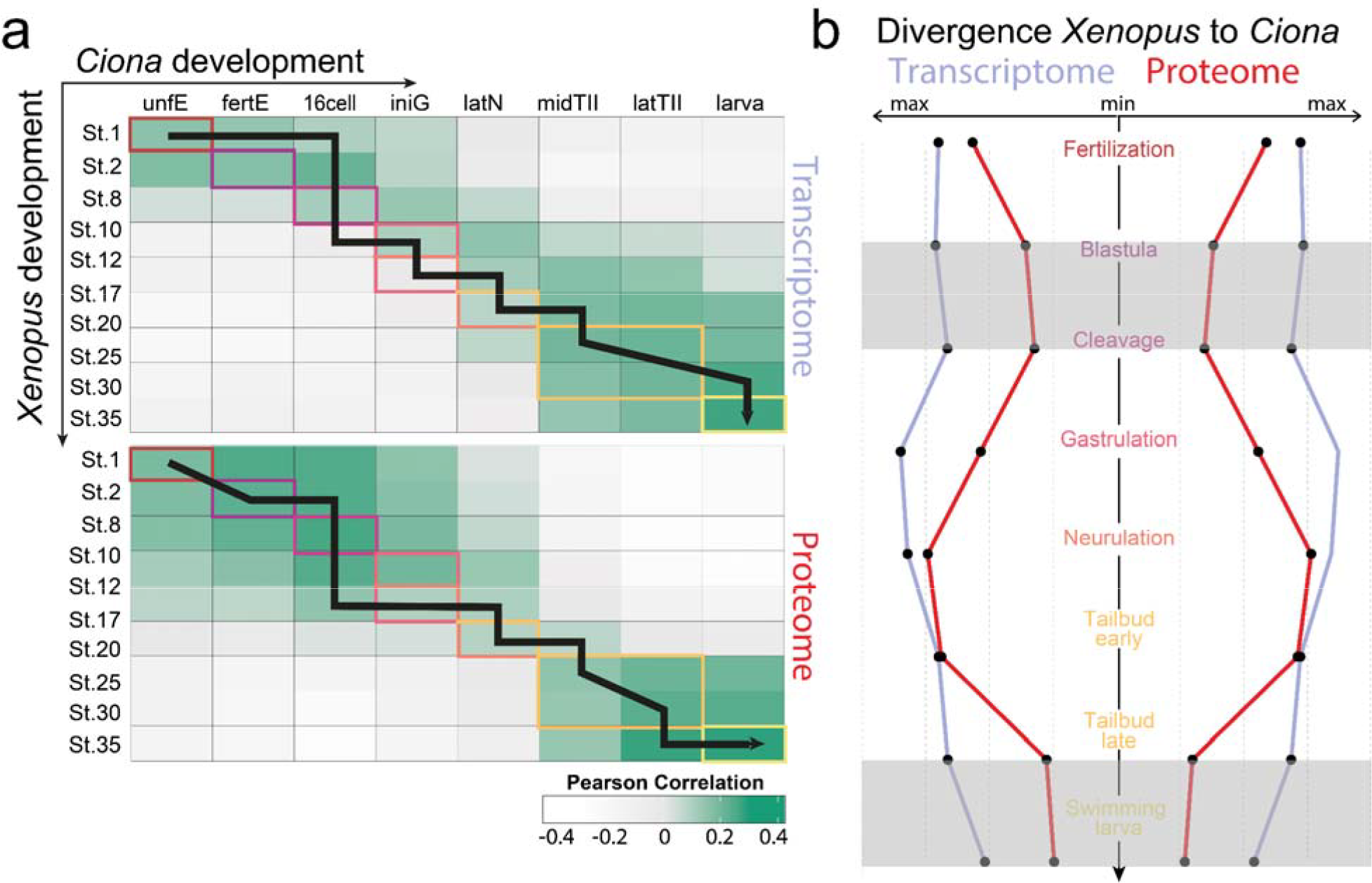
The protein anti-hourglass model. a) Similarity heatmaps showing Pearson similarity between the two species for each investigated time point. Developmental stages are color-coded as defined in a. The black line follows the highest correlation of the *Xenopus* time-point for each *Ciona* stage. b) Temporal divergence of gene and protein expression at individual developmental stages during embryogenesis between *Ciona* and *Xenopus*. Maximal similarity is represented by the smallest distance from the center line, revealing a nested hourglass model in which the proteome exhibits more evident bottlenecks at early and later stages. Gray boxes outline these periods of minimal divergence. Regardless of stage, proteins show higher similarity between the two species’ developmental mapping than RNA-seq, suggesting that protein dynamics are evolutionary more conserved than mRNA dynamics.

When we plot the highest correlation of mRNA and protein expression for each stage of *Ciona*, we find that for each stage, protein dynamics are more highly correlated than corresponding mRNA dynamics (Fig. 4b). This suggests that protein dynamics are evolutionarily more conserved than mRNA dynamics likely due to proteins conveying function (Laurent et al. 2010b; Schrimpf et al. 2009b). It seems likely that post-transcriptional mechanisms, e.g. varying translation or protein degradation rates, have evolved to compensate for different mRNA dynamics.

We find that mRNA and protein expression profiles are comparatively diverse in unfertilized eggs, but converge during cleavage stages and then exhibit maximum divergence at either gastrulation (via RNA-seq) or neurulation (via proteomics) (Kourakis et al. 2021). We hypothesize that this divergence might represent the distinct mechanisms of gastrulation and neurulation in the two species. In *Ciona*, gastrulation takes place via a cup-shaped gastrula driven by invagination of the endoderm, whereas in *Xenopus*, convergent extension of mesoderm and epidermal epiboly play important roles. Most importantly, *Ciona* differs temporally from its vertebrate cousin by specifying its axis at the neurula stage, rather than at gastrulation (Winkley et al. 2019).

Together, our data both on the proteome and transcriptome provide evidence for an anti-hourglass model, whereby gene activity is most divergent at mid-developmental stages (Hu et al. 2017b; Levin et al. 2016b; Marlétaz et al. 2018b) (Fig 4b). The embryos show most similarity at the larval stage, likely due to common structural and ecological requirements in swimming larvae (Fig 4 b). As evidenced by the anti-hourglass pattern of expression, this method readily captures evolutionary changes within the chordate phylum. Strikingly, the anti-hourglass is much more pronounced on the proteome than the transcriptome level (Fig. 4b) demonstrating the potential of using proteomics for evolutionary cross-species comparison. Our evolutionary analysis underscores the significance of the simple chordate *Ciona* as a linchpin in unraveling conserved aspects of chordate and vertebrate embryogenesis, particularly in future comparative studies.

In this study on the simple chordate *Ciona*, we provide protein estimates in the unfertilized egg as well as a rich multi-omic atlas of transcriptome and proteome dynamics throughout embryogenesis. To our knowledge this represents the deepest proteomic dataset and an excellent starting point to further explore the intricate dynamics of proteogenomics. We additionally generated a cross-species time-series comparison between *Ciona* and *Xenopus* highlighting similarities and differences in mRNA and protein expression. We show that the proteome correlates more strongly throughout all compared developmental stages. Both our proteomics and transcriptomics comparisons provide evidence for the inverse hourglass model suggesting that *Ciona* and *Xenopus* are maximally diverse at mid-developmental stages. We believe that such comparative studies are greatly served by the use of integrated transcriptomic and proteomic datasets rather than relying solely on RNA-seq comparisons.

## Materials and Methods

### SNP prevalence between *Ciona* batches

One concern is the presence of single nucleotide polymorphisms (SNPs), a characteristic feature of ascidian evolution (Abdul-Wajid et al. 2014; Tsagkogeorga et al. 2012b), which can cause protein sequence polymorphisms and lead to incorrect peptide inference during the processing of MS data. We evaluated the potential influence of SNPs on peptide quantification accuracy. We obtained bulk RNA-seq data from two batches of 16-cell *Ciona* embryos. Each batch was assembled via Trinity, then translated into protein reference databases with the Mass Spec Protein Reference tool (https://kirschner.med.harvard.edu/tools/mz_ref_db.html) (Wühr et al. 2014). We reciprocally BLASTed each database against the other and found 16,037 shared proteins. These shared proteins were Trypsin digested *in silico*. 98.8% of the resulting peptides were identical between these batches while only 1.2% were wholly unique to one batch or the other indicating minimal influence of intra-specific genetic variability on peptide recognition.

### Generating protein reference database

Assembly of the protein reference database -a fasta file containing all potential proteins from the species under study used to generate *in silico* tryptic peptides and reference MS/MS spectra for peptide identification -was done mostly as described previously (Wühr et al. 2014). 1,222,451,669 *Ciona* bulk RNA-seq reads from numerous studies (Reeves et al. 2017; Wang et al. 2019; Kaplan et al. 2019; Sharma et al. 2019) were assembled *de novo* via Trinity (version 2.11) (Grabherr et al. 2011) into 2,328,005 transcripts. The 55,974 transcripts making up the KH *Ciona* transcriptome (KHNCBI.Transcript.2018.fasta, retrieved from ANISEED) (Brozovic et al. 2016) were integrated alongside our *de novo* transcripts. The transcripts were cleaned and trimmed via SeqClean (http://compbio.dfci.harvard.edu/tgi/software/), then masked for common repeat motifs via RepeatMasker (version 4.1) (Smit et al. 2015). The masked transcripts were clustered via TGICL (version 2.1) and assembled via CAP3 (Huang and Madan 1999) using parameters discussed previously. The resulting contigs and singletons were searched against a database of model organism containing human (*Homo sapiens*), Red junglefowl (*Gallus gallus*), Western clawed frog (*Xenopus tropicalis*), zebrafish (*Danio rerio*), Florida lancelet (*Branchiostoma floridae*), Pacific purple sea urchin (*Strongylocentrotus purpuratus*), and urochordate (*Ciona robusta*) using BLASTX (version 2.10.1) (Altschul et al. 1990). The BLASTX report was parsed as described previously to translate the transcripts into proteins. The translated proteins were processed as described previously, however with a CD-HIT (version 4.8.1) (Li and Godzik 2006; Fu et al. 2012) threshold of 95%.

### *Ciona* handling and embryos collection

Wild type adult *Ciona robusta* (formerly known as *Ciona intestinalis* Type A) (Brunetti et al. 2015) were obtained from M-Rep located in San Diego, CA and maintained in artificial seawater (Instant Ocean) at 18 °C, under continuous illumination. Dechorionation and *in vitro* fertilization procedures were conducted following the protocol described in (Christiaen et al. 2009). For each time point in the time series, embryos were staged and collected according to (Hotta et al. 2007) at approximately 18 °C and a total of 150 embryos were placed in Trizol for RNA extraction, while approximately 3000 embryos were rapidly frozen in liquid nitrogen for protein TMTproC sample preparation. All samples were then stored at -80 °C until further use. For absolute mass spectrometry analysis, approximately 5000 unfertilized eggs were directly snap-frozen.

### Proteomics sample preparation

Samples were prepared by lysing frozen embryos in lysis buffer (50 mM HEPES pH 7.2, 2% SDS, and 1x protease in artificial saltwater) followed by clarification via centrifugation. Lysates were diluted to 2 ug/μL with 100 mM HEPES (pH 7.2). DTT was added to a concentration of 5 mM and samples incubated for 20 mins at 60 °C. After cooling to RT, N-ethylmaleimide (NEM) was added to a concentration of 20 mM and samples incubated for 20 mins at RT. 10 mM DTT was added and samples incubated for 10 mins at RT to quench NEM. To 200 μL of each sample were brought up to 2 mL with 800 μL MeOH, 400 μL chloroform, and 600 μL water. Samples were centrifuged at 20,000 g for 2 minutes at RT. Upper layer was discarded and 600 μL MeOH was added. Samples were centrifuged at 20,000 g for 2 minutes at RT. Supernatant was discarded and 500 μL MeOH was added. Samples were centrifuged at 20,000 g for 2 minutes at RT. Supernatant was discarded and the pellet was air dried. Pellet was resuspended in 6 M GuaCl, 10 mM EPPS pH 8.5 to ∼5 mg/mL.

For the label-free samples, UPS2 standards (Sigma-Aldrich) were added to a final concentration of 27 ng/μL in the 450 μg protein samples. Samples were diluted with 10 mM EPPS pH 8.5 to 2 M guanidine hydrochloride. Samples were digested overnight at RT in LysC (Wako) at a concentration of 20 ng/μL. Samples were further diluted with 10 mM EPPS pH 8.5 to 0.5 M guanidine hydrochloride. 20 ng/μL LysC and 10 ng/μL Trypsin (Promega) were added to each sample and incubated for 16 hours at 37 °C. Peptide supernatant was cleared by ultracentrifugation at 100,000□g for 1 hour at 4□°C (Beckman Coulter, 343775), then vacuum-dried overnight.

For TMTpro-labeling, samples were digested with LysC and Trypsin as above, then resuspended in 200 mM EPPS pH 8.0. pre-mixed TMTpro tags (8-plex Thermo Fisher Scientific 20 μg/μL in dry acetonitrile stored at -80 °C) at a 5 μg TMTpro: 1 μg peptide ratio. Tags are as follows: 126 -unfertilized egg; 128C – fertilized egg; 129N – 16-cell; 130C – 110-cell; 131N – late neurula; 131C – mid tailbud II; 133C – late tailbud II; 134N – larva. Samples were incubated for 2 hours at RT. Reactions were quenched by addition of hydroxylamine (Sigma, HPLC grade) to a final concentration of 0.5% for 30 minutes at RT. Samples were pooled into a single tube, cleared by ultracentrifugation at 100,000 g for 1 hour at 4□°C (Beckman Coulter, 343775), then and vacuum-dried overnight.

For either label-free or TMTpro-labeled, samples were resuspended with 10□mM ammonium bicarbonate (pH 8.0) with 5% acetonitrile to 1 μg/μL. Samples were separated by medium pH reverse phase HPLC (Zorbax 300Extend C18, 4.6 x 250 mm column) into 96 fractions as described previously (Johnson et al. 2021a; Nguyen et al. 2022). The fractions were then pooled into 24 fractions (Edwards and Haas 2015), dried, and resuspended in HPLC grade water. Samples were then desalted via homemade stage tips with C18 material (Empore) and resuspended to 1 μg/μL in 1% formic acid.

### Proteomics analysis

Approximately 1□μg per sample was analyzed by LC-MS, as previously described (Nguyen et al. 2022). LC-MS experiments were analyzed on an nLC-1200 HPLC (Thermo Fisher Scientific) coupled to an Orbitrap Fusion Lumos MS (Thermo Fisher Scientific). Peptides were separated on an Aurora Series emitter column (25□cm□×□75□μm ID, 1.6□μm C18) (Ionopticks, Australia), held at 60□°C during separation by an in-house built column oven. Separation was achieved by applying a 12 % to 35 % acetonitrile gradient in 0.125 % formic acid and 2 % DMSO over 90□minutes for fractionated samples. Electrospray ionization was enabled by applying a voltage of 2.6□kV through a MicroTee at the inlet of the microcapillary column. For the TMTpro samples, we used the Orbitrap Fusion Lumos with the TMTproC method previously described (Johnson et al. 2021a). For the label-free samples, we used the Orbitrap Fusion Lumos with the label-free method previously described (Wühr et al. 2014).

Mass spectrometry data analysis was performed essentially as previously described (Sonnett et al. 2018b) with the following modifications. The raw MS files were analyzed using the Gygi Lab software platform (GFY Core) licensed through Harvard University. MS2 spectra assignment was performed using the SEQUEST algorithm by searching the data against either our reference protein dataset described above, the KY21 *Ciona* proteome (Satou et al. 2021), or the Uniprot *Ciona* proteome. To control for peptide false discovery rate, reverse sequences were searched in parallel with forward sequences (Elias and Gygi 2007). For label-free analysis, these proteomes were merged with the UPS2 Proteomics Standards FASTA file (Sigma-Aldrich) along with common contaminants. Peptides that matched multiple proteins were assigned to the proteins with the greatest number of unique peptides. TMTproC data were analyzed as previously described (Johnson et al. 2021a).

### Proteomics analysis Absolute protein concentration estimates

Protein concentration in the egg was calculated by building a standard curve of MS signal to UPS2 standard concentration. The UPS2 known standard concentrations were obtained from Sigma Aldrich and concentrations were converted to log space. The MS signal area was also converted to log space and thiel regression was performed to obtain a standard curve. Signal area for *Ciona* egg protein analyzed with the KY21 proteome (Satou et al. 2021) was converted to concentration and scaled to a total protein concentration of 2 mM. A cutoff of 0.01 μM was applied for low concentration protein.

### RNA Sequencing

A total of 150 embryos from each developmental stage were collected and stored at -80 °C in Trizol. Total RNA was isolated using the Clean and Concentrator Zymo kit (Zymo), with genomic DNA (gDNA) removal achieved through on-column treatment with Turbo DNase (Invitrogen) at room temperature for 10 minutes. The resulting RNA was re-suspended in 15 μl of DEPC-treated water and quantify using a NanodropTM and Qubit (Thermo Fisher Scientific, Waltham, USA), while its quality was assessed using a Bioanalyzer 2100 (Agilent Technologies, Santa Clara, USA). The RNA integrity number (RIN) values ranged between 8 and 10. cDNA libraries were prepared using the PrepX RNA-seq directional protocol (Takara Bio) following the manufacturer’s instructions and utilizing an Apollo 324 robot. For mRNA enrichment and separation from rRNA, the oligo dT-based mRNA isolation kit (Takara Bio) was employed. The libraries were sequenced on the NovaSeq platform (Illumina, San Diego, USA) with a depth of 20-40 million reads per sample.

Quality assessment of the raw and trimmed 61 bp paired RNA-seq reads was performed using FastQC. Trimgalore (version 0.6.10) was used to trim the raw RNA-seq reads, removing adapter contamination and poor quality base calls. The trimmed RNA-seq reads were then mapped to the KY21 transcriptome using Salmon (version 0.43.1), and subsequent downstream analysis were conducted in R and Python.

### *Ciona* and *Xenopus* protein orthologs and pairwise comparison

Standard protein-protein BLAST (BLASTP, version 2.10.1) was used to identify orthologs between *Ciona* and *Xenopus (Altschul et al. 1990). Ciona* and *Xenopus* alternated as query and reference. For each BLASTP, the max target sequence was set to 1, e-value threshold was set to 0.01, and the matrix set to BLOSUM45. The query ID, reference ID, e-value, and bit score were logged for each match. A python script was used to identify the “best-match” protein orthologs between *Ciona* and *Xenopus* based on the criteria of (1) lowest e-value and (2) highest bit score. Only proteins confirmed in both directions as “best-match” were used in the cross-species proteomic analysis. Similarity between *Ciona* and frog stages was measured by pairwise Pearson’s correlation based on one-to-one orthologs identified using OrthoFinder (v.2.5.4) (Emms and Kelly 2019).

### Drawings

*Ciona* schematics are adapted from FABA (FABA Four-dimensional Ascidian Body Atlas) and *Xenopus* illustrations form Xenbase © Natalya Zahn (2022).

## Acknowledgements

We thank all members of the Wühr laboratory for helpful discussion, particularly to Felix Keber and Edward Cruz. We also thank members of the Levine laboratory, especially Pavan Choppakatla for insightful inputs. We thank Nicholas Treen for providing unfertilized *Ciona* eggs and Lillia Ryazanova for assistance in *Ciona* protein sample preparation.

## Author contributions

ANF, AM, MSL and MW conceived the project. ANF, AM, MSL and MW designed the experiments. ANF and AM performed the experiments and data analysis. All authors interpreted results and drafted the manuscript. MSL and MW supervised the project.

## Data and materials availability

All data needed to evaluate the conclusions in the paper will be available on Github upon publication. The raw sequencing data and gene expression matrices are available in GEO under the accession number: X. The mass spectrometry proteomics data have been deposited to the ProteomeXchange Consortium via the PRIDE partner repository with the dataset identifier X.

## Bibliography

Abdulghani M, Song G, Kaur H, Walley JW, Tuteja G. 2019. Comparative Analysis of the Transcriptome and Proteome during Mouse Placental Development. J Proteome Res 18: 2088–2099.

Abdul-Wajid S, Veeman MT, Chiba S, Turner TL, Smith WC. 2014. Exploiting the extraordinary genetic polymorphism of ciona for developmental genetics with whole genome sequencing. Genetics 197: 49–59.

Abitua PB, Gainous TB, Kaczmarczyk AN, Winchell CJ, Hudson C, Kamata K, Nakagawa M, Tsuda M, Kusakabe TG, Levine M. 2015. The pre-vertebrate origins of neurogenic placodes. Nature 524: 462–465.

Abitua PB, Wagner E, Navarrete IA, Levine M. 2012. Identification of a rudimentary neural crest in a non-vertebrate chordate. Nature 492: 104–107.

Ahn HR, Kim GJ. 2012. The Ascidian Numb Gene Involves in the Formation of Neural Tissues. Dev Reprod 16: 371–378.

Altschul SF, Gish W, Miller W, Myers EW, Lipman DJ. 1990. Basic local alignment search tool. J Mol Biology 215: 403–410.

Ammar C, Schessner JP, Willems S, Michaelis AC, Mann M. 2023. Accurate label-free quantification by directLFQ to compare unlimited numbers of proteomes. Mol Cell Proteom 100581.

Baxi AB, Lombard-Banek C, Moody SA, Nemes P. 2018. Proteomic Characterization of the Neural Ectoderm Fated Cell Clones in the Xenopus laevis Embryo by High-Resolution Mass Spectrometry. ACS Chem Neurosci 9: 2064–2073.

Baxi AB, Pade LR, Nemes P. 2021. Mass spectrometry based proteomics for developmental neurobiology in the amphibian Xenopus laevis. Curr Top Dev Biology 145: 205–231.

Berthelot C, Villar D, Horvath JE, Odom DT, Flicek P. 2018. Complexity and conservation of regulatory landscapes underlie evolutionary resilience of mammalian gene expression. Nat Ecol Evol 2: 152–163.

Brozovic M, Martin C, Dantec C, Dauga D, Mendez M, Simion P, Percher M, Laporte B, Scornavacca C, Di Gregorio A, et al. 2016. ANISEED 2015: a digital framework for the comparative developmental biology of ascidians. Nucleic Acids Res 44: D808–D818.

Brunetti R, Gissi C, Pennati R, Caicci F, Gasparini F, Manni L. 2015. Morphological evidence that the molecularly determined Ciona intestinalis type A and type B are different species: Ciona robusta and Ciona intestinalis. J Zoo□l Syst Evol Res 53: 186–193.

Bultman SJ, Gebuhr TC, Pan H, Svoboda P, Schultz RM, Magnuson T. 2006. Maternal BRG1 regulates zygotic genome activation in the mouse. Genes Dev 20: 1744–1754.

Cao C, Lemaire LA, Wang W, Yoon PH, Choi YA, Parsons LR, Matese JC, Wang W, Levine M, Chen K. 2019. Comprehensive single-cell transcriptome lineages of a proto-vertebrate. Nature 571: 349–354.

Chan ME, Bhamidipati PS, Goldsby HJ, Hintze A, Hofmann HA, Young RL. 2021. Comparative Transcriptomics Reveals Distinct Patterns of Gene Expression Conservation through Vertebrate Embryogenesis. Genome Biol Evol 13: evab160.

Christiaen L, Wagner E, Shi W, Levine M. 2009. Isolation of sea squirt (Ciona) gametes, fertilization, dechorionation, and development. Cold Spring Harb Protoc 2009: pdb.prot5344.

Christmas MJ, Kaplow IM, Genereux DP, Dong MX, Hughes GM, Li X, Sullivan PF, Hindle AG, Andrews G, Armstrong JC, et al. 2023. Evolutionary constraint and innovation across hundreds of placental mammals. Science 380: eabn3943.

Dehal P, Satou Y, Campbell RK, Chapman J, Degnan B, Tomaso AD, Davidson B, Gregorio AD, Gelpke M, Goodstein DM, et al. 2002. The Draft Genome of Ciona intestinalis: Insights into Chordate and Vertebrate Origins. Science 298: 2157–2167.

Delsuc F, Brinkmann H, Chourrout D, Philippe H. 2006. Tunicates and not cephalochordates are the closest living relatives of vertebrates. Nature 439: 965–968.

Delsuc F, Philippe H, Tsagkogeorga G, Simion P, Tilak M-K, Turon X, López-Legentil S, Piette J, Lemaire P, Douzery EJP. 2018. A phylogenomic framework and timescale for comparative studies of tunicates. BMC Biol 16: 39.

Demichev V, Messner CB, Vernardis SI, Lilley KS, Ralser M. 2020. DIA-NN: neural networks and interference correction enable deep proteome coverage in high throughput. Nat Methods 17: 41–44.

Edfors F, Danielsson F, Hallström BM, Käll L, Lundberg E, Pontén F, Forsström B, Uhlén M. 2016. Gene□specific correlation of RNA and protein levels in human cells and tissues. Mol Syst Biol 12: 883.

Edwards A, Haas W. 2015. Multiplexed Quantitative Proteomics for High-Throughput Comprehensive Proteome Comparisons of Human Cell Lines. Methods Mol biology Clifton N J 1394: 1–13.

Elias JE, Gygi SP. 2007. Target-decoy search strategy for increased confidence in large-scale protein identifications by mass spectrometry. Nat Methods 4: 207–214.

Emms DM, Kelly S. 2019. OrthoFinder: phylogenetic orthology inference for comparative genomics. Genome Biol 20: 238.

Evans VC, Barker G, Heesom KJ, Fan J, Bessant C, Matthews DA. 2012. De novo derivation of proteomes from transcriptomes for transcript and protein identification. Nat Methods 9: 1207–1211.

Fu L, Niu B, Zhu Z, Wu S, Li W. 2012. CD-HIT: accelerated for clustering the next-generation sequencing data. Bioinformatics 28: 3150–3152.

Ghaemmaghami S, Huh W-K, Bower K, Howson RW, Belle A, Dephoure N, O’Shea EK, Weissman JS. 2003. Global analysis of protein expression in yeast. Nature 425: 737–741.

Ghazalpour A, Bennett B, Petyuk VA, Orozco L, Hagopian R, Mungrue IN, Farber CR, Sinsheimer J, Kang HM, Furlotte N, et al. 2011. Comparative Analysis of Proteome and Transcriptome Variation in Mouse. PLoS Genet 7: e1001393.

Gil-Gálvez A, Jiménez-Gancedo S, Pérez-Posada A, Franke M, Acemel RD, Lin C-Y, Chou C, Su Y-H, Yu J-K, Bertrand S, et al. 2022. Gain of gene regulatory network interconnectivity at the origin of vertebrates. Proc National Acad Sci 119: e2114802119.

Goldberger R. 1980. Biological Regulation and Development, Molecular Organization and Cell Function.

Grabherr MG, Haas BJ, Yassour M, Levin JZ, Thompson DA, Amit I, Adiconis X, Fan L, Raychowdhury R, Zeng Q, et al. 2011. Trinity: reconstructing a full-length transcriptome without a genome from RNA-Seq data. Nat Biotechnol 29: 644–652.

Gupta M, Sonnett M, Ryazanova L, Presler M, Wühr M. 2018. Quantitative Proteomics of Xenopus Embryos I, Sample Preparation. Methods Mol Biology Clifton N J 1865: 175–194.

Haeckel E. 1866. bd. Allgemeine entwickelungsgeschichte der organismen. G. Reimer.

Horie R, Hazbun A, Chen K, Cao C, Levine M, Horie T. 2018. Shared evolutionary origin of vertebrate neural crest and cranial placodes. Nature 560: 228–232.

Hotta K, Mitsuhara K, Takahashi H, Inaba K, Oka K, Gojobori T, Ikeo K. 2007. A web □based interactive developmental table for the ascidian Ciona intestinalis, including 3D real□image embryo reconstructions: I. From fertilized egg to hatching larva. Dev Dyn 236: 1790–1805.

Hu H, Uesaka M, Guo S, Shimai K, Lu T-M, Li F, Fujimoto S, Ishikawa M, Liu S, Sasagawa Y, et al. 2017a. Constrained vertebrate evolution by pleiotropic genes. Nat Ecol Evol 1: 1722–1730.

Huang X, Madan A. 1999. CAP3: A DNA Sequence Assembly Program. Genome Res 9: 868–877.

Imai KS, Hino K, Yagi K, Satoh N, Satou Y. 2004a. Gene expression profiles of transcription factors and signaling molecules in the ascidian embryo: towards a comprehensive understanding of gene networks. Development 131: 4047–4058.

Imai KS, Levine M, Satoh N, Satou Y. 2006. Regulatory Blueprint for a Chordate Embryo. Science 312: 1183–1187.

Izzi SA, Colantuono BJ, Sullivan K, Khare P, Meedel TH. 2013. Functional studies of the Ciona intestinalis myogenic regulatory factor reveal conserved features of chordate myogenesis. Dev Biology 376: 213–223.

Johnson A, Stadlmeier M, Wu□hr M. 2021a. TMTpro Complementary Ion Quantification Increases Plexing and Sensitivity for Accurate Multiplexed Proteomics at the MS2 Level. J Proteome Res 20: 3043–3052.

Kaplan NA, Wang W, Christiaen L. 2019. Initial characterization of Wnt-Tcf functions during Ciona heart development. Dev Biology 448: 199–209.

Keller TE, Han P, Yi SV. 2016. Evolutionary Transition of Promoter and Gene Body DNA Methylation across Invertebrate–Vertebrate Boundary. Mol Biol Evol 33: 1019–1028.

Kourakis MJ, Bostwick M, Zabriskie A, Smith WC. 2021. Disruption of left-right axis specification in Ciona induces molecular, cellular, and functional defects in asymmetric brain structures. BMC Biol 19: 141.

Kubo A, Suzuki N, Yuan X, Nakai K, Satoh N, Imai KS, Satou Y. 2010. Genomic cis-regulatory networks in the early Ciona intestinalis embryo. Development 137: 1613–1623.

Laurent JM, Vogel C, Kwon T, Craig SA, Boutz DR, Huse HK, Nozue K, Walia H, Whiteley M, Ronald PC, et al. 2010a. Protein abundances are more conserved than mRNA abundances across diverse taxa. PROTEOMICS 10: 4209–4212.

Laurent JM, Vogel C, Kwon T, Craig SA, Boutz DR, Huse HK, Nozue K, Walia H, Whiteley M, Levin M, Anavy L, Cole AG, Winter E, Mostov N, Khair S, Senderovich N, Kovalev E, Silver DH, Feder M, et al. 2016a. The mid-developmental transition and the evolution of animal body plans. Nature 531: 637–641.

Li W, Godzik A. 2006. Cd-hit: a fast program for clustering and comparing large sets of protein or nucleotide sequences. Bioinformatics 22: 1658–1659.

Lopez CE, Sheehan HC, Vierra DA, Azzinaro PA, Meedel TH, Howlett NG, Irvine SQ. 2017. Proteomic responses to elevated ocean temperature in ovaries of the ascidian Ciona intestinalis. Biol Open 6: 943–955.

Madgwick A, Magri MS, Dantec C, Gailly D, Fiuza U-M, Guignard L, Hettinger S, Gomez-Skarmeta JL, Lemaire P. 2019. Evolution of embryonic cis-regulatory landscapes between divergent Phallusia and Ciona ascidians. Dev Biol 448: 71–87.

Marlétaz F, Firbas PN, Maeso I, Tena JJ, Bogdanovic O, Perry M, Wyatt CDR, Calle-Mustienes E de la Bertrand S, Burguera D, et al. 2018a. Amphioxus functional genomics and the origins of vertebrate gene regulation. Nature 564: 64–70.

McAlister GC, Nusinow DP, Jedrychowski MP, Wu□hr M, Huttlin EL, Erickson BK, Rad R, Haas W, Gygi SP. 2014. MultiNotch MS3 Enables Accurate, Sensitive, and Multiplexed Detection of Differential Expression across Cancer Cell Line Proteomes. Anal Chem 86: 7150–7158.

Messerschmidt DM, Vries W de, Ito M, Solter D, Ferguson-Smith A, Knowles BB. 2012. Trim28 Is Required for Epigenetic Stability During Mouse Oocyte to Embryo Transition. Science 335: 1499–1502.

Nguyen T, Costa EJ, Deibert T, Reyes J, Keber FC, Tomschik M, Stadlmeier M, Gupta M, Kumar CK, Cruz ER, et al. 2022. Differential nuclear import sets the timing of protein access to the embryonic genome. Nat Commun 13: 5887.

Nomura M, Nakajima A, Inaba K. 2009a. Proteomic profiles of embryonic development in the ascidian Ciona intestinalis. Dev Biol 325: 468–481.

Pappireddi N, Martin L, Wühr M. 2019. A Review on Quantitative Multiplexed Proteomics. ChemBioChem 20: 1210–1224.

Peshkin L, Wühr M, Pearl E, Haas W, Freeman RM, Gerhart JC, Klein AM, Horb M, Gygi SP, Kirschner MW. 2015a. On the Relationship of Protein and mRNA Dynamics in Vertebrate Embryonic Development. Dev Cell 35: 383–394.

Reeves WM, Wu Y, Harder MJ, Veeman MT. 2017. Functional and evolutionary insights from the Ciona notochord transcriptome. Development 144: 3375–3387.

Santesmasses D, Mariotti M, Guigó R. 2017. Selenoproteins, Methods and Protocols. Methods Mol Biology 1661: 17–28.

Satou Y, Hamaguchi M, Takeuchi K, Hastings KEM, Satoh N. 2006. Genomic overview of mRNA 5′-leader trans-splicing in the ascidian Ciona intestinalis. Nucleic Acids Res 34: 3378–3388.

Satou Y, Nakamura R, Yu D, Yoshida R, Hamada M, Fujie M, Hisata K, Takeda H, Satoh N. 2019. A Nearly Complete Genome of Ciona intestinalis Type A (C. robusta) Reveals the Contribution of Inversion to Chromosomal Evolution in the Genus Ciona. Genome Biol Evol 11: 3144–3157.

Satou Y, Tokuoka M, Oda-Ishii I, Tokuhiro S, Ishida T, Liu B, Iwamura Y. 2022. A Manually Curated Gene Model Set for an Ascidian, Ciona robusta (Ciona intestinalis Type A). Zool Sci 39: 253–260.

Saxena S, Dupont S, Meghah V, Lakshmi MGM, Singh SK, Swamy CVB, Idris MM. 2013. Proteome map of the neural complex of the tunicate Ciona intestinalis, the closest living relative to vertebrates. PROTEOMICS 13: 860–865.

Schrimpf SP, Weiss M, Reiter L, Ahrens CH, Jovanovic M, Malmström J, Brunner E, Mohanty S, Lercher MJ, Hunziker PE, et al. 2009a. Comparative Functional Analysis of the Caenorhabditis elegans and Drosophila melanogaster Proteomes. PLoS Biol 7: e1000048.

Schwanhäusser B, Busse D, Li N, Dittmar G, Schuchhardt J, Wolf J, Chen W, Selbach M. 2011. Global quantification of mammalian gene expression control. Nature 473: 337–342.

Session AM, Uno Y, Kwon T, Chapman JA, Toyoda A, Takahashi S, Fukui A, Hikosaka A, Suzuki A, Kondo M, et al. 2016. Genome evolution in the allotetraploid frog Xenopus laevis. Nature 538: 336–343.

Sharma S, Wang W, Stolfi A. 2019. Single-cell transcriptome profiling of the Ciona larval brain. Dev Biology 448: 226–236.

Sladitschek HL, Fiuza U-M, Pavlinic D, Benes V, Hufnagel L, Neveu PA. 2020. MorphoSeq: Full Single-Cell Transcriptome Dynamics Up to Gastrulation in a Chordate. Cell 181: 922–935.e21.

Smit, Hubley, Green R&. 2015. RepeatMasker Open-4.0. http://www.repeatmasker.org(Accessed 2015).

Smits AH, Lindeboom RGH, Perino M, Heeringen SJ van Veenstra GJC, Vermeulen M. 2014a. Global absolute quantification reveals tight regulation of protein expression in single Xenopus eggs. Nucleic Acids Res 42: 9880–9891.

Sonnett M, Gupta M, Nguyen T, Wühr M. 2018a. Quantitative Proteomics for Xenopus Embryos II, Data Analysis. Methods Mol Biology Clifton N J 1865: 195–215.

Sonnett M, Yeung E, Wu □hr M. 2018b. Accurate, Sensitive, and Precise Multiplexed Proteomics Using the Complement Reporter Ion Cluster. Anal Chem 90: 5032–5039.

Stolfi A, Gainous TB, Young JJ, Mori A, Levine M, Christiaen L. 2010. Early Chordate Origins of the Vertebrate Second Heart Field. Science 329: 565–568.

Stolfi A, Ryan K, Meinertzhagen IA, Christiaen L. 2015. Migratory neuronal progenitors arise from the neural plate borders in tunicates. Nature 527: 371–374.

Sun L, Champion MM, Huber PW, Dovichi NJ. 2016. Proteomics of Xenopus development. MHR Basic Sci reproductive medicine 22: 193–199.

Suzuki MM, Mori T, Satoh N. 2016. The Ciona intestinalis cleavage clock is independent of DNA methylation. Genomics 108: 168–176.

Thompson A, Schäfer J, Kuhn K, Kienle S, Schwarz J, Schmidt G, Neumann T, Johnstone R, Mohammed AKA, Hamon C. 2003. Tandem Mass Tags: A Novel Quantification Strategy for Comparative Analysis of Complex Protein Mixtures by MS/MS.Anal Chem 75: 1895–1904.

Touceda-Suárez M, Kita EM, Acemel RD, Firbas PN, Magri MS, Naranjo S, Tena JJ, Gómez-Skarmeta JL, Maeso I, Irimia M. 2020. Ancient genomic regulatory blocks are a source for regulatory gene deserts in vertebrates after whole genome duplications. Mol Biol Evol 37: msaa123..

Tsagkogeorga G, Cahais V, Galtier N. 2012a. The Population Genomics of a Fast Evolver: High Levels of Diversity, Functional Constraint, and Molecular Adaptation in the Tunicate Ciona intestinalis. Genome Biol Evol 4: 852–861.

Tsitsiridis G, Steinkamp R, Giurgiu M, Brauner B, Fobo G, Frishman G, Montrone C, Ruepp A. 2022. CORUM: the comprehensive resource of mammalian protein complexes–2022. Nucleic Acids Res 51: D539–D545.

Uesaka M, Kuratani S, Irie N. 2022. The developmental hourglass model and recapitulation: An attempt to integrate the two models. J Exp Zoo □l Part B, Mol Dev Evol 338: 76–86.

Veeman MT, Nakatani Y, Hendrickson C, Ericson V, Lin C, Smith WC. 2007. chongmague reveals an essential role for laminin-mediated boundary formation in chordate convergence and extension movements. Development 135: 33–41.

Vogel C, Marcotte EM. 2012. Insights into the regulation of protein abundance from proteomic and transcriptomic analyses. Nat Rev Genet 13: 227–232.

Walton KD, Croce JC, Glenn TD, Wu S-Y, McClay DR. 2006. Genomics and expression profiles of the Hedgehog and Notch signaling pathways in sea urchin development. Dev Biol 300: 153–164.

Wan L-B, Pan H, Hannenhalli S, Cheng Y, Ma J, Fedoriw A, Lobanenkov V, Latham KE, Schultz RM, Bartolomei MS. 2008. Maternal depletion of CTCF reveals multiple functions during oocyte and preimplantation embryo development. Development 135: 2729–2738.

Wang W, Niu X, Stuart T, Jullian E, Mauck WM, Kelly RG, Satija R, Christiaen L. 2019. A single-cell transcriptional roadmap for cardiopharyngeal fate diversification. Nat Cell Biology 21: 674–686.

Wegler C, Ölander M, Wiśniewski JR, Lundquist P, Zettl K, Ásberg A, Hjelmesæth J, Andersson TB, Artursson P. 2019. Global variability analysis of mRNA and protein concentrations across and within human tissues. Nar Genom Bioinform 2: qz010.

Winkley KM, Kourakis MJ, DeTomaso AW, Veeman MT, Smith WC. 2019. Tunicate gastrulation. Curr Top Dev Biol 136: 219–242.

Wühr M, Freeman RM, Presler M, Horb ME, Peshkin L, Gygi SP, Kirschner MW. 2014. Deep Proteomics of the Xenopus laevis Egg using an mRNA-Derived Reference Database. Curr Biol 24: 1467–1475.

Yamada L, Saito T, Taniguchi H, Sawada H, Harada Y. 2009. Comprehensive Egg Coat Proteome of the Ascidian Ciona intestinalis Reveals Gamete Recognition Molecules Involved in Self-sterility*. J Biol Chem 284: 9402–9410.

Zhang T, Xu Y, Imai K, Fei T, Wang G, Dong B, Yu T, Satou Y, Shi W, Bao Z. 2020. A single-cell analysis of the molecular lineage of chordate embryogenesis. Sci Adv 6: eabc4773.

