## Supplemental figures for "Quantitative proteome dynamics across embryogenesis in a model chordate"

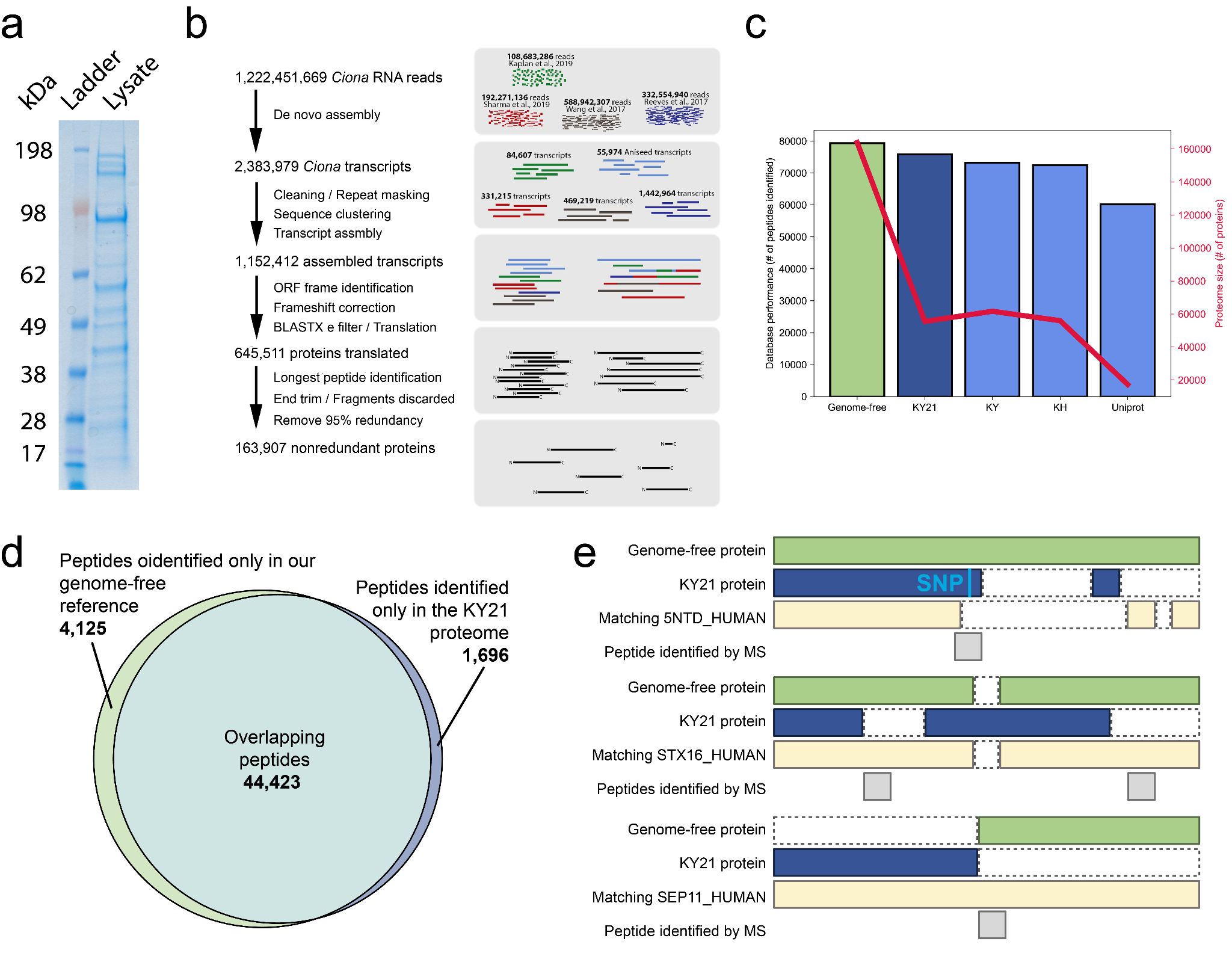

**Supplemental Fig 1: Summary of yolk abundance and genome-free protein reference database.**

a) *Ciona* egg lysate analysis with SDS-PAGE gel and coomassie blue stain. *Ladder:* SeeBlue Plus2 pre-stained protein standard. *Lysate:* Total *Ciona* egg lysate. Strong signal for the 100 kDa apolipoprotein-B-like yolk protein (Yamada et al., 2009) was observed.

b) Schematics of assembly. 1,222,451,669 RNA-seq reads were aggregated from 5 different studies and translated into a protein reference database as described previously (Wühr et al., 2014) to obtain 163,907 95% non-redundant proteins.

c) The genome-free reference database, and several *Ciona* proteomes, were used to analyze the same TMTproC MS dataset. The bar heights correspond to the total number of peptides identified in the MS dataset for each annotation. The red line represents the size of the proteome database. Our reference database outperforms the KY21 proteome slightly in terms of peptide identification, but at the expense of a much larger size and manual annotation of proteins.

d) Comparison of peptides identification using genome-free reference database and KY21 proteome reference. The Venn diagram illustrates the shared and unique peptides identified using each reference.

e) Examples where genome-free reference can help improve current KY21 assembly from SNPs detection (blue insert), correction of mis-annotated coding sequences, and accurate annotation of selenoproteins.

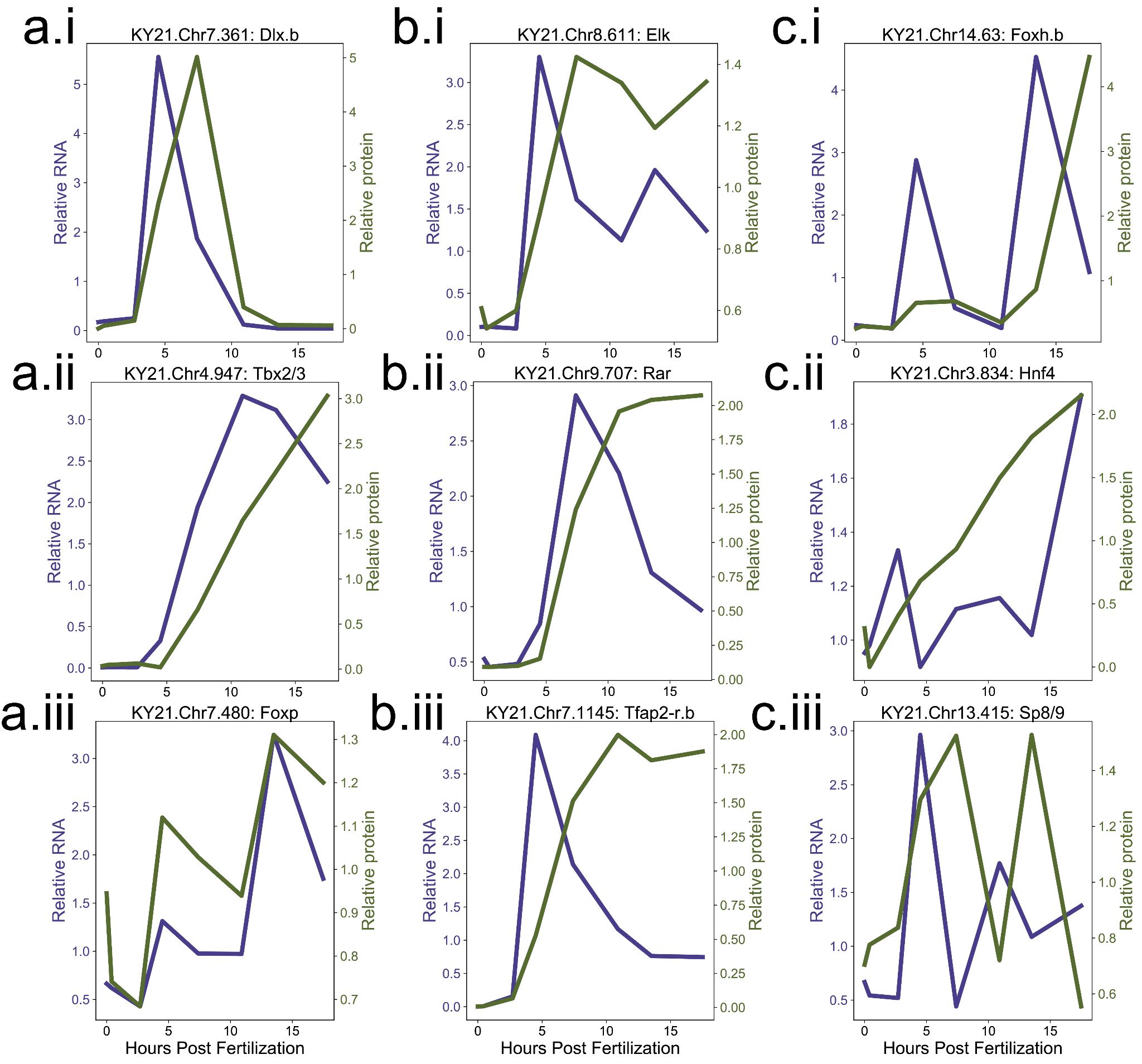

**Supplemental Fig 2: Different RNA and protein dynamics during development for TFs with known involvement in early *Ciona* development.**

a.i-iii) Dlx.b, Tbx2/3, and Foxp have similar RNA and protein dynamics, largely matching in relative expression across development.

b.i-iii) Elk, Rar, and Tfap2-r.b have similar trends in their respective RNA and protein dynamics, with RNA being expressed earlier than protein and degrading while protein expression remains high.

c.i-iii) Foxy.b, Hnf4, and Sp8/9 do not have strong trends in their RNA and protein dynamics. RNA and protein expression seem more sporadic, with RNA coming in distinct waves that are not necessarily followed by protein.
